# Bit-parallel sequence-to-graph alignment

**DOI:** 10.1101/323063

**Authors:** Mikko Rautiainen, Veli Mäkinen, Tobias Marschall

## Abstract

Graphs are commonly used to represent sets of sequences. Either edges or nodes can be labeled by sequences, so that each path in the graph spells a concatenated sequence. Examples include graphs to represent genome assemblies, such as string graphs and de Bruijn graphs, and graphs to represent a pan-genome and hence the genetic variation present in a population. Being able to align sequencing reads to such graphs is a key step for many analyses and its applications include genome assembly, read error correction, and variant calling with respect to a variation graph. Here, we generalize two linear sequence-to-sequence algorithms to graphs: the *Shift-And* algorithm for exact matching and Myers’ *bitvector* algorithm for semi-global alignment. These linear algorithms are both based on processing *w* sequence characters with a constant number of operations, where *w* is the word size of the machine (commonly 64), and achieve a speedup of *w* over naive algorithms. Our bitvector-based graph alignment algorithm reaches a worst case runtime of 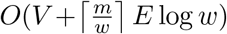 for acyclic graphs and *O*(*V* + *mE* log *w*) for arbitrary cyclic graphs. We apply it to four different types of graphs and observe a speedup between 3.1-fold and 10.1-fold compared to previous algorithms.

## 1 Introduction

Aligning two sequences is a classic problem in bioinformatics. The standard dynamic programming (DP) algorithm, introduced by Needleman and Wunsch in 1970 [15], aligns two sequences of length *n* in *O*(*n*^2^) time. Countless variants of this classic DP algorithm exist, in particular its generalization to local alignment [20], where the alignment can be between any substrings of the two sequences, and semi-global alignment [19] where one sequence (*query*) is entirely aligned to a substring of the other (*reference*).

In addition to sequences, graphs whose nodes or edges are labeled by characters are commonly used for many applications in bioinformatics, for instance for genome assembly [11, 4] and multiple sequence alignment [7]. Currently, we witness a strong interest in the use of graphs also as a potential alternative to “linear” reference genomes [5, 17]. Graph-based reference genomes hold the promise of removing reference biases and can naturally encode also complex variation. With an increasing usage of graphs, algorithms for aligning reads to graphs are also of growing interest and have already been applied successfully for purposes such as genome assembly [1] and error correction [18]. So far, however, algorithms to align sequences to graphs while exploiting bit-parallelism have been lacking.

**Related Work** In this paper, we study the semi-global sequence-to-graph alignment problem. That is, we seek to find a path in the directed, node-labeled graph that has minimum edit distance to the query sequence. Already in 1989, an algorithm was discovered for approximate regular expression matching [12], which represented the regular expression as a graph and achieved a runtime of *O*(*V* + *mE*) for aligning a sequence to it. In 2000, an *O*(*V* + *mE*) algorithm for aligning a sequence to an arbitry graph was discovered in the context of hypertext searching [14]. The algorithm is a generalization of the Needleman-Wunsch algorithm for graphs. It proceeds row-wise with two sweeps per row: on the first sweep, calculating the recurrence from the values in the previous row, and on the second sweep, propagating the recurrence term for the values in the same row with a depth first search.

Other algorithms for sequence-to-graph alignment have been discovered in the context of bioinformatics; however, although published later than the *O*(*V* + *mE*) algorithms, they either obtain worse runtimes, do not apply to arbitrary graphs, or do not produce the optimal alignment. We list these results below for completeness. *Partial order alignment* [8] (POA) extends standard DP to directed acyclic graphs (DAG) in *O*(*V* + *mE*) time but does not handle cyclic graphs. The variation graph tool vg [6] aligns to cyclic graphs by “unrolling” the graph into a DAG, and then uses POA. However, unrolling the graph can produce a drastically larger DAG [22]. V-align [22] aligns to arbitrary graphs with *O*((*V*′ + 1)*mE*) runtime where V’ is the graph’s minimum feedback vertex set. Limasset et al. align reads to de Bruijn graphs [9], but in a heuristic manner without guaranteeing optimal alignment. The genome assembler hybridSPAdes [1] re-phrases sequence-to-graph alignment as a shortest path problem and uses Dijkstra’s algorithm, leading to *O*(|*E*|*m* + |*V*|*m*log(|*V*|*m*)) runtime.

**Contributions** In this paper, we introduce techniques for bit-parallel semi-global sequence-to-graph alignment. To illustrate some of the central ideas, we first discuss the simpler question of generalizing the *Shift-And* algorithm [2] for exact string matching to graphs. We obtain an algorithm with an 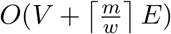 runtime in acyclic graphs, matching the Shift-And algorithm for linear sequences, and *O*(*V* + *mE*) runtime in arbitrary cyclic graphs. We then generalize Myers’ *bitvector* alignment algorithm [13] to graphs, which proceeds along the same lines as for the Shift-And algorithm, but requires some further algorithmic insights to merge bit-vectors representing columns of the alignment matrix. We arrive at an algorithm with a runtime of 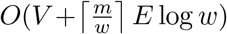 for acyclic graphs and *O*(*V* + *mE* log *w*) for arbitrary cyclic graphs. Moreover, we perform experiments showing that despite the higher time complexity in cyclic graphs, the bitvector algorithm is empirically faster than the *O*(*V* + *mE*) algorithm for hypertext searching [14] by a factor of 3.1 to 10.1, depending on the input graphs.

## 2 Problem definition

### Definition 1

(Sequence graph). We define a *sequence graph* as a tuple *G* = (*V*, *E*, *σ*), where *V* = [*υ*,…, *υ*_*n*_} is a finite set of nodes, *E* ⊂ *V* × *V* is a set of directed edges and *σ* : *V* → Σ assigns one character from the alphabet Σ to each node. We refer to the sets of indices of in-neighbors and out-neighbors of node *υ*_*i*_ as 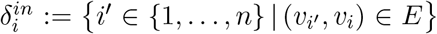 and 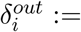 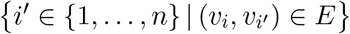, respectively.

### Definition 2

(Path sequence). Let *p* = (*p*_1_,…,*p*_*k*_) be a path in the sequence graph *G* = (*V*, *E*, *σ*); that is, *p*_*i*_ ∈ *V* for *i* ∈ {1,…, *k*} and (*p*_*i*_,*p*_*i*_+_1_) ∈ *E* for *i* ∈ {1,…, *k* − 1}. Then, the *path sequence* of *p*, written *σ*(*p*), is given by *σ*(*p*_1_)*σ*(*p*_2_) ⃛ *σ*(*p*_*k*_).

We note that this definition of paths and path sequences includes the possibility of repeated vertices. In this paper, we study two related graph problems: finding exact matches between a sequence and a path in a graph and the semi-global sequence-to-graph alignment (SG^2^A) problem.

### Problem 1

(Sequence-to-Graph Exact Matching). Let a string *s* ∈ Σ* and a sequence graph *G* = (*V*, *E*, *σ*) be given. Find all paths *p* = (*p*_1_,&, *p*_*k*_) in *G* such that the path label *σ*(*p*) is equal to the string *s*, or report that such a path does not exist.

### Problem 2

(Unit Cost Semi-Global Sequence-to-Graph Alignment). Let a string *s* ∈ Σ* and a sequence graph *G* = (*V*, *E*, *σ*) be given. Find a path *p* = (*p*_*i*_,…, *p*_*k*_) in *G* such that the edit distance *d*(*σ*(*p*),*s*) is minimized and report a corresponding alignment of *σ*(*p*) and *s*.

Note that the exact matching problem is easier in the sense that Problem 2 can be used to solve Problem 1 by finding paths with an edit distance of 0. In the remainder of this paper, we assume an arbitrary but fixed string *s* ∈ Σ* with |*s*| = *m* and sequence graph *G* = (*V*,*E*,*σ*) to be given. Without loss of generality, we assume that |*V*| ∈ *O*(|*E*|). If |*V*| > 2|*E*|, there are nodes which are not connected to any other nodes. We can merge the disconnected nodes with the same label, producing a graph with 2|*E*| + |Σ| nodes at most.

## 3 Extending Shift-And to graphs

The *Shift-And* algorithm [2, 3] finds exact matches between a pattern string *s* of size *m* and a text string *t* of size *n*, with *m* < *n*, in 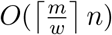 time where *w* is the word size of the machine (64). The Shift-And algorithm works by simulating a nondeterministic finite automaton (NFA) that matches the pattern, and then feeding the text to it. The state of the automaton is kept in a *m*-sized bitvector, consisting of 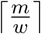 *w*-bit words, and the state is updated by shifting the vector by one and AND-ing the state with a precomputed character bitvector. The invariant of the algorithm is that the *i*’th bit in the NFA’s state is set after processing the *j*’th character in the text if and only if there is an exact match between the pattern at *s*_0․.*i*_ and the text at *t*_*j*−*i*‥*j*_.

### 3.1 DAGs

In directed acyclic graphs (DAG), we order the nodes topologically and then process them in topological order. However, some nodes have an in-degree of more than 1. For handling nodes with an in-degree more than 1, we first propagate the NFA state from each in-neighbor separately. Since the path to the node can come from any of the in-neighbors, we need to merge the states such that any exact match from any in-neighbor translates to a match in the node. The invariant of the algorithm is that a set bit means an exact match. As the match can come from any in-neighbor, this means we have to merge the states with bitwise OR. Since the merging is a 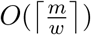-time operation, the time complexity is unchanged.

### 3.2 Cyclic graphs

The strategy for cyclic regions is similar to the previous, except that, in the absence of a topological sorting, we process the edges in an arbitrary order. The main idea to still arrive at correct values consists in storing a separate NFA state bit-vector for each graph node and to update them repeatedly until no more changes are necessary.

Algorithm 1 shows our algorithm as pseudocode. We keep a list of *calculable* edges. All edges are inserted into the calculable list at the start. Whenever a node’s NFA state has changed, we add all outgoing edges to the list. A state change may set a bit but it cannot unset a bit. Therefore a node’s state may change up to m times, so each edge may get added to the list up to m times. The worst case runtime is therefore *O*(*V* + *mE*). Correctness can be verified by observing that the above invariant must hold for all nodes once the calculable list is empty. Algorithm 1 can be simplified to the 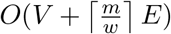 algorithm for DAGs by sorting *L* topologically, popping the edges in order at line 7, and removing the IF block starting from line 10. For the DAG algorithm, we also do not need to keep the entire array *S*, but just a “frontier” consisting of nodes whose out-neighbors have not been processed yet.

## 4 Extending Myers’ bitvector alignment to graphs

We approach Problem 2 by generalizing the standard dynamic programming (DP) algorithm for edit distance calculation. In our case, the DP matrix has one column per node *υ*_*i*_ ∈ *V* and one row per character *s_j_* from *s* ∈ Σ*. We seek to compute values *C*_*i,j*_ for *i* ∈ {1,…, |*V*|} and *j* ∈ {1,…, |*s*|} such that *C*_*i,j*_ is the minimum edit distance *d*(*p*, *s*[1‥*j*]) over all paths *p* ending in node υ_*i*_.

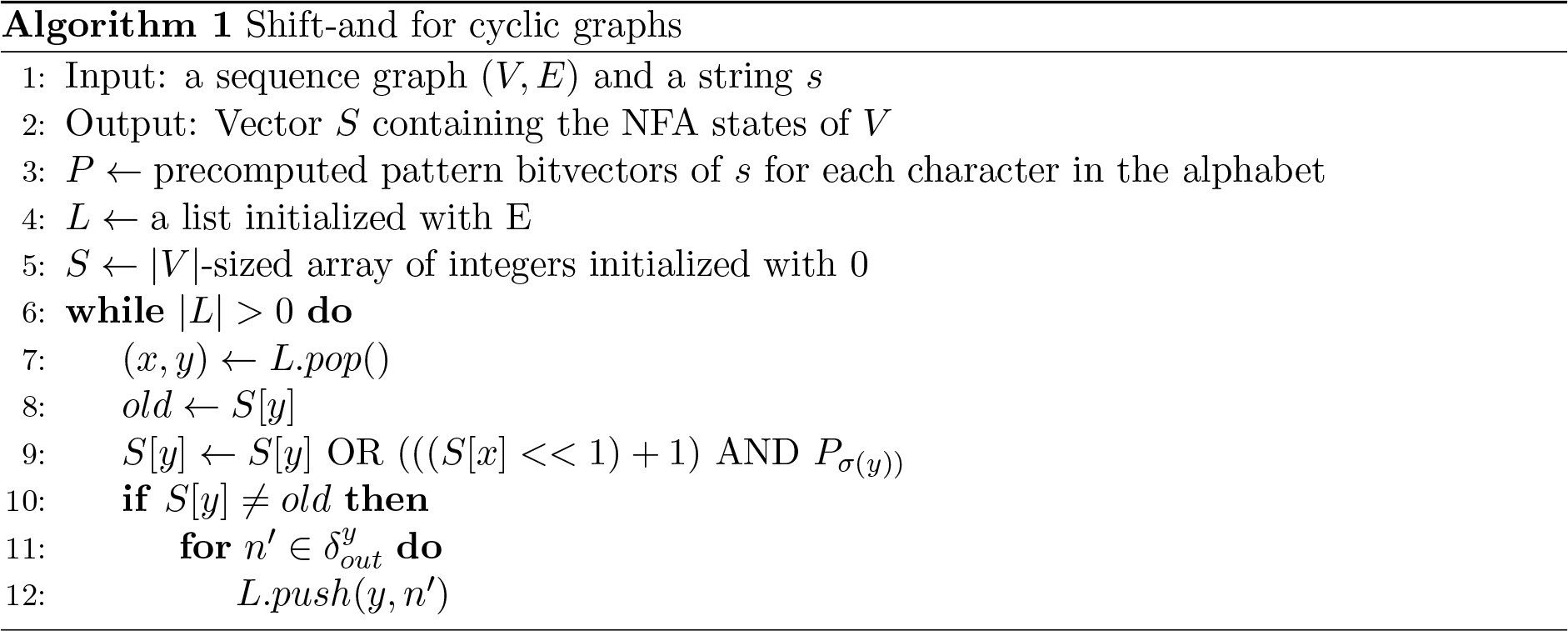

**Figure 1:**
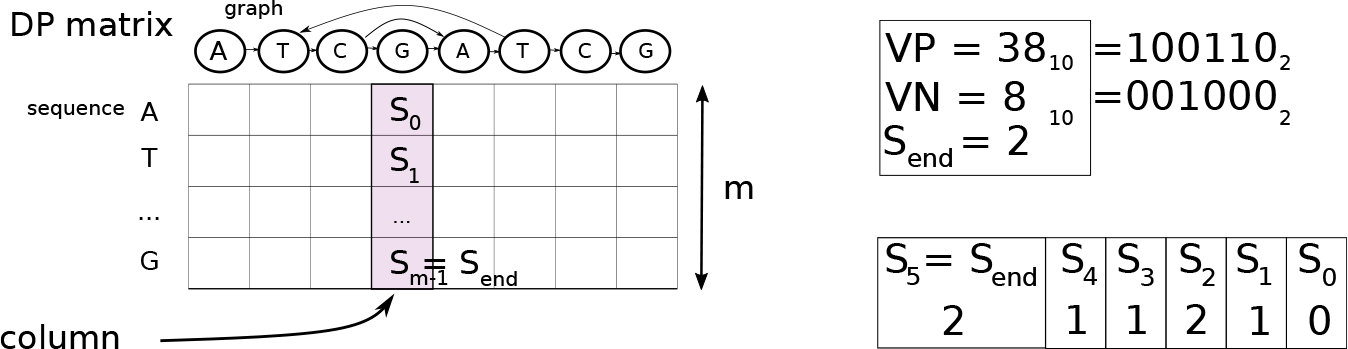
Left: The relations between the DP matrix (grid), columns (purple rectangle), and *m*. Right: Explicit representation of a bitvector (upper rectangle) and the implicit scores (lower rectangle), with *m* = 6.

### Definition 3

(Recurrence for SG^2^A). Set *C*_*i*,1_ = Δ_*i*,1_ for all *i* ∈ {1,…, |*V*|}. And define 
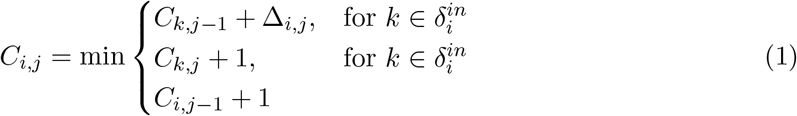
where Δ_*i,j*_ the mismatch penalty between node character *σ*(*υ*_*i*_) and sequence character *s*_*j*_, which is 0 for a match and 1 for a mismatch.

Note that despite the cyclic dependencies in Recurrence (1), the problem has a unique solution for any graph and sequence. Recurrence (1) can be solved in *O*(*V* + *mE*) time [14] in a *cell-by-cell* manner, where each operation calculates one individual cell. This is in contrast to Myers’ bitvector algorithm for sequence-to-sequence alignment which calculates multiple cells in a constant time operation [13].

In linear sequence-to-sequence alignment, the recurrence implies the *vertical property* [21], meaning that the score difference between two vertically neighboring cells is in the range {−1,0,1}, which is necessary for representing them using two *bitvectors* [13]. To generalize Myers’ algorithm, we first establish that the vertical property also holds for graphs.

### Theorem 1

(Vertical property for sequence-to-graph alignment). The score difference between any two vertically adjacent cells *C*_*i,j*_ and *C*_*i,j−1*_ is at most one, that is, *C*_*i,j*_ − *C*_*i,j−1*_ ∈ {−1,0,1} for all *i* ∈ {1,…, |*V*|} and *j* ∈ {2,…, |*s*|}.

*Proof.* See Appendix.

### 4.1 Terminology

Figure 1 shows the relation between the concepts described here. The DP matrix is oriented with graph characters as columns and sequence characters as rows. A *column* is an *m*-cell vertical column in the DP matrix, corresponding to one node in the graph. We use the terms *column* and *node* interchangeably, depending on if we are emphasizing the DP matrix or the graph topology. We use the term *calculating a column* to refer to the operation of using Recurrence (1) to calculate a destination column from a source column and a character (the label of destination node/column). The *minimum changed score* between two columns *C*^old^ and *C*^new^ is the minimum score of the *new* column at rows where the new column is smaller, that is, 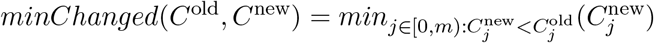. If 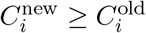 at every *i*, we say that the minimum changed score is infinite. The minimum changed score is used to distinguish cells which are relevant in cyclic areas; when recalculating a column, only those cells whose scores changed can propagate the scores onward.

**Figure 2:**
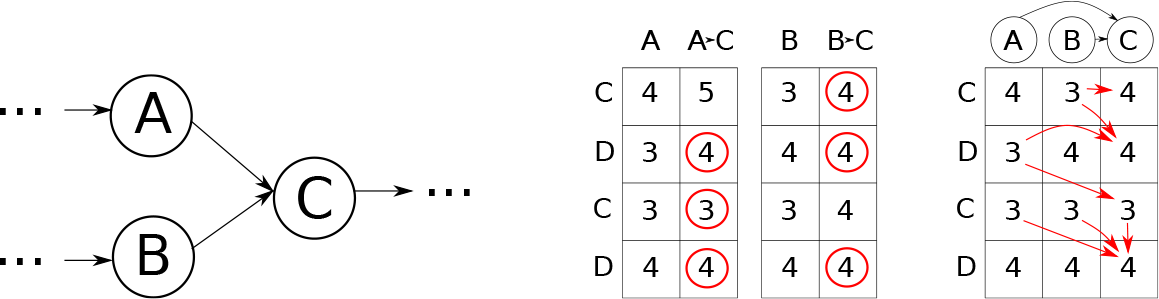
Handling nodes with an in-degree higher than one in the bitvector framework. Left: The node C has two in-neighbors, A and B. Middle: Each in-neighbor column is separately calculated to get the scores of Recurrence (1). The circled cells are the minimum of each row. Right: The resulting column are merged, taking the minimum of the two scores for each row. The arrows show where each cell got its value from.

We refer to the current DP table column we consider as *S*. Column *S* is stored in *bitvector representation* [13], consisting of a score *S*_*end*_ attained in that column at the bottom row, a positive bitvector *VP*^*S*^ and a negative bitvector *VN*^*S*^. The *word size w* is the number of bits in a computer word (usually 64). The positive and negative bitvectors consist of *m* bits and are implemented with 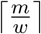 machine words. For a column *S*, the *score* at index *i* is 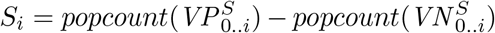, where *popcount* refers to the number of set bits in a bitvector. Note that *S*_*end*_ = *S*_*m−1*_.

### 4.2 Directed acyclic graphs

For DAGs, we use a similar strategy to the Shift-And algorithm. First we order the nodes topologically, and then we calculate the columns in order. However, Recurrence (1) now has terms for multiple in-neighbors. For handling nodes with an in-degree more than 1, we first calculate the incoming column from each in-neighbor, that is, as if there was only one in-neighbor. Then, we merge the columns such that the cells of the resulting column have the minimum of each incoming column in that row. Figure 2 shows an example of this. For merging more than two columns, we simply merge them two at a time: denoting the column merge operator as ⨂, the result of merging columns *C*_1_,*C*_2_,*C*_3_,… is *C*_*r*_ = ((*C*_1_ ⨂ *C*_2_) ⨂ *C*_3_)…. We defer the details of merging columns to Section 5, where we devise an algorithm to do this in 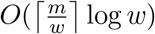 time. The operation must be applied at most *E* times. The runtime is therefore 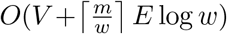.

### 4.3 Cyclic regions

Cell-by-cell algorithms for sequence-to-graph alignment [14, 12] handle cyclic dependencies in a row-wise manner: For each row, in a first sweep the “vertical” and “diagonal” terms of Recurrence (1) are calculated and, in a second sweep, the “horizontal” terms are applied. However, this approach cannot be applied in a column-wise manner that is inherent to Myers’ bitvector algorithm. To deal with cyclic dependencies, we rely on two key ideas: First, we process the edges in a specific order and, second, we recalculate scores of nodes until they have “converged” (similar to our approach for the Shift-And algorithm).

The define this order, we keep a queue of *calculable edges* and their priorities. Initially, all edges are inserted into the queue with priority 0. All columns are initialized with a bitvector *VP* = 1^*m*^, *VN* = 0^*m*^, corresponding to increasing scores. Then, edges are picked from the queue in priority order (lowest first), and the target column is calculated based on the source column. The new column is merged with the existing column and the merged column is stored at the node. Then, if the minimum changed value between the existing and the new column is not infinite, the target node’s out-neighbors are added to the calculable queue with the minimum changed value as the priority. Pseudocode is given in Algorithm 2. We use the symbol ⨂ to mark the column merging operation. We use the *F* to denote the column calculation operation from a predecessor column and a character match bitvector. This operation proceeds exactly like in Myers’ original bitvector algorithm and involves computing intermediate bitvectors for horizontal and diagonal differences. We do not discuss these details here and refer the reader to the original paper [13] or to the textbook [10]. In the following, we will establish correctness and runtime of Algorithm 2.

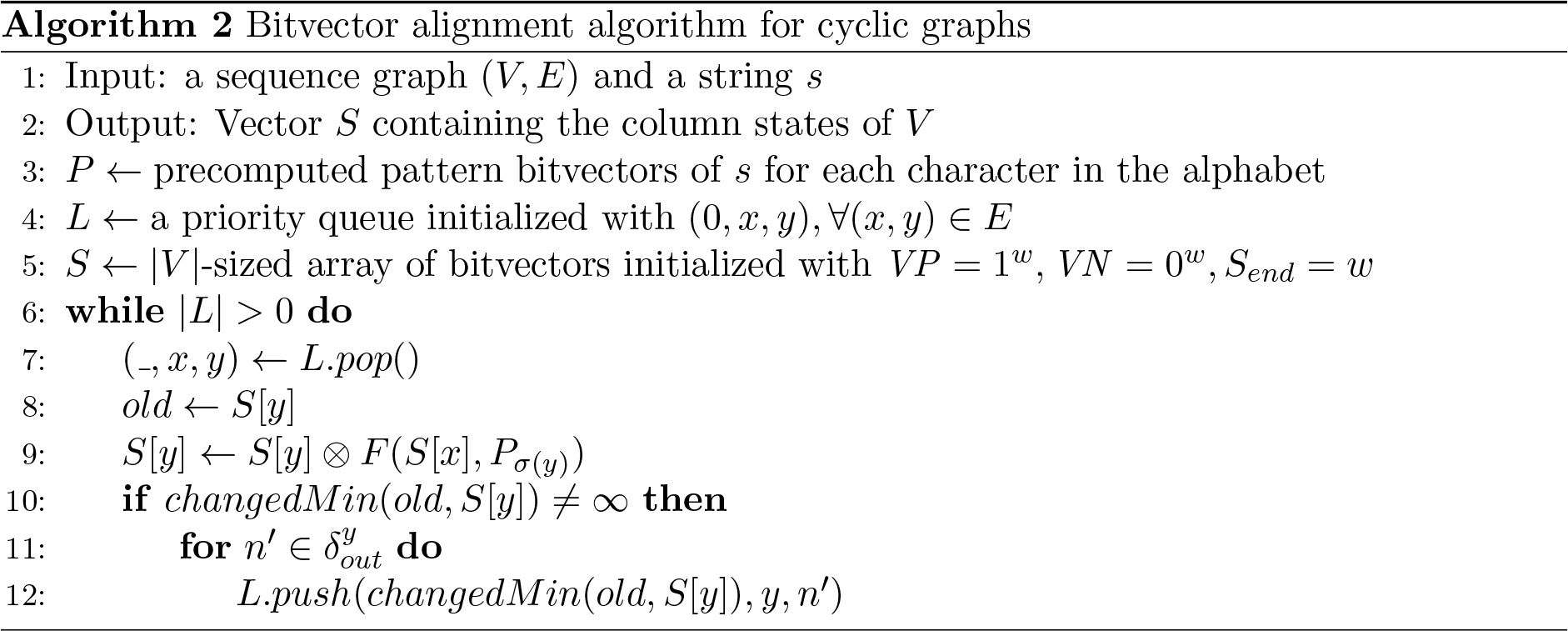

We use the term *present scores* to refer to the scores assigned to the cells at some point during the calculation, as opposed to the *correct scores* which correspond to the unique scores that satisfy Recurrence (1). We say that a cell has *converged* when its present score is equal to its correct score.

#### Theorem 2.

In Algorithm 2, if the minimum priority of the calculable queue is *x*, then all cells whose correct scores are *C*_*i,j*_ < *x* have converged.

*Proof*. We show this by induction. For the initial case, there are no cells whose correct scores are negative, so the statement holds when *x* = 0. Next, we will assume that the minimum priority of the calculable queue is *x* and that all cells whose correct scores are *C*_*i,j*_ < *x* − 1 have converged, and show that all cells whose correct scores are *C*_*i,j*_ = *x* − 1 have converged. Assume that there is a cell whose correct score is *x* − 1. There are four cases for how the cell’s correct score is defined: (i) the vertical term, (ii) the horizontal term, (iii) the diagonal term with a mismatch, (iv) the diagonal neighbor with a match.

*Case (i)*. The cell has a vertical neighbor *C*_*i,j*−1_ whose correct score is *x* − 2. By assumption cells with correct score *C*_*i*′,*j*′_ < *x* − 1 have converged, so the vertical neighbor’s present score is *x* − 2. The bitvector representation allows a vertical score difference of up to 1, so the cell’s present score is at most *x* − 1 and the cell has converged.

*Case (ii)*. The cell has a horizontal neighbor *C*_**i*′,*j**_ whose correct score is x − 2. The neighbor cell has converged by assumption. After the last time the neighbor column was calculated, the neighbor cell had its correct score. Since there is a cell with a present score *x* − 2 in the neighboring column, the edge (*i′*, *i*) was added to the calculable queue with a priority of *x* − 2 (or less). Therefore the edge (*i′*, *i*) was processed at some point earlier in the calculation, and at that point Recurrence (1) was applied to the cell *C*_*i,j*_, producing the correct score.

*Case (iii)*. Analogous to Case (ii).

*Case (iv)*. The cell has a diagonal neighbor *C*_*i′,j′*_ whose correct score is *x* − 1. If the diagonal neighbor has converged, then the edge (*i′*, *i*) will have been added to the calculable queue with a priority of *x* − 1 (or less), and the argument from case (ii) applies. Next we need to prove that the diagonal neighbor has converged. The diagonal neighbor cell’s correct score is again defined by the same cases (i)-(iv). For cases (i)-(iii), the diagonal neighbor has converged. For case (iv), we look at the diagonal neighbor cell’s diagonal neighbor cell, and keep traversing by diagonal connections until we reach a cell for whom one of cases (i)-(iii) applies. Since the diagonal neighbors cannot form cycles, this will eventually happen, proving that the entire chain has converged.

From Theorem 2 it follows that once the minimum priority of the calculable queue is *m* + 1, all cells have converged to their correct scores, so the algorithm will eventually reach the correct solution in cyclic areas. Next we will establish an upper bound on the time until convergence.

#### Lemma 3.

For an edge with priority *x* in the calculable queue, applying Recurrence (1) from the source node to the target node produces at least one cell with a present score of at most *x* + 1.

*Proof*. The edge’s priority is given by its source column’s minimum changed score, which is the present score at some cell in the column. Since the edge has a priority *x*, at least one cell in the source column has a present score of *x*. From Recurrence (1), the horizontal and diagonal neighbor cells of that cell receive a present score of at most *x* + 1.

#### Corollary 4.

If all cells whose correct scores are *C*_*i,j*_ < *x* have converged, then all cells whose present scores are *C*_*i,j*_ ≤ *x* have converged.

*Proof.* We assumed that all cells whose correct scores are *C*_*i,j*_ < *x* have converged. Therefore there are no cells whose present score is *x* but whose correct score is *C*_*i,j*_ < *x*. A cell’s present score cannot be lower than its correct score since the present scores are initialized at the highest possible value and applying Recurrence (1) cannot lower them under the correct score. Therefore if a cell’s present score is *x*, it must also be its correct score.

#### Theorem 5.

An edge cannot be added to the calculable queue more than 2*m* times

*Proof.* Edges are added to the calculable queue whenever their source column’s present scores change. From Lemma 3, an edge with a priority *x* produces at least one cell with a present score of at most *x* + 1. If the score is *x* or less, then the cell has converged by Theorem 2 and Corollary 4. In that case, the cell’s present score cannot change anymore, so this can happen up to m times. If the cell has a present score of *x* + 1, then the cell may or may not have converged. If it has converged, the cell’s present score cannot change anymore. If it does not have converged, then the cell’s present score can change one more time later, for a total of 2 changes per cell. Since there are *m* cells in a column, the column’s state can change at most 2*m* times.

Theorem 5 provides a bound for the running time: an edge cannot be processed more than *m* times, meaning that in the worst case, the cyclic bitvector algorithm behaves like a cell-by-cell algorithm. In Section 5, we will see how partitioning the DP matrix in horizontal slices of *w* rows results in an asymptotically faster algorithm.

Calculating a column produces a column with scores at most one higher than the source column. This means that the changed minimum value between the target node’s old and new columns can be at most *w* + 1 higher than the priority of the source node. Therefore the priority of the newly added edges is at most *w* + 1 higher than the minimum priority in the calculable queue. By using *w* + 1 arrays as the calculable queue and swapping pointers when the current queue is empty, inserting and retrieving *n* values can be done in *O*(*m* + *n*) time where m is the largest value which can be inserted, in this case the length of the sequence. Since *n* > *m*, this reduces to *O*(*n*) and the calculable queue has in practice amortized constant time retrieval and insertion.

In summary, calculating one edge takes 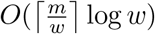 time due to merging the columns and calculating the changed minimum value. This means that the total runtime of Algorithm 2 is 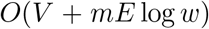. Like the Shift-And algorithm, the cyclic algorithm can also be simplified for DAGs by ordering; *L* topologically in line 4 and removing the IF-block starting at line 10, producing an 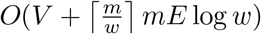 algorithm.

## 5 Bitvector implementation

The scores of each column are represented with a *bitvector* consisting of a positive bitvector *VP*, negative bitvector *VN* and score at end *S*_*end*_. For a sequence of length *m*, the bitvectors consist of *m* bits, implemented as 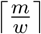 machine words. The bitvectors can be further sliced into smaller parts, and the size of the parts affects the runtime. We use the term *elementary operation* to refer to arithmetic and bitwise operations, and the term *column operation* to refer to the bitvector merging and minimum changed score algorithms described here. For a bitvector of *k* bits, the column operations use *O*(log *k*) elementary operations. For *m*-bit bitvectors, the elementary operations are 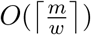, meaning that the column operations are 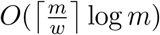. However, if we slice the bitvector into w-bit slices, elementary operations are *O*(1) and column operations *O*(log *w*) per slice. With 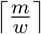 slices, this leads to a total runtime of 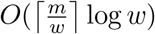 for elementary operations and 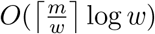 for column operations. Due to the bitvector slicing, we also have to calculate the DP matrix in 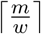 horizontal slices, each consisting of *w* rows. The algorithm described in the previous section is applied to each *w*-row slice in order from top to bottom. With the slicing, the bitvectors also include an extra term *score before start S*_*before*_, which is 0 for the topmost slice and equal to the above slice’s *S*′_*end*_ for other slices.

### 5.1 Bitvector merging algorithm

Figure 3 contains a running example of merging two bitvectors with an *O*(log *w*) algorithm, which we use to explain the algorithm. For completeness, we provide full pseudocode in the Appendix as Algorithm 3 and refer to the respective line numbers below.

The input for the bitvector merging algorithm are two bitvectors *A* and *B*, consisting of values *VP*^*A*^, *VN*^*A*^, 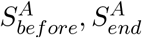 and *VP*^*B*^, *VN*^*B*^, 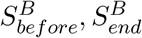(step A). We assume that 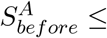 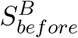. These bitvectors implicitly represent the values 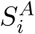 and 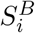 (step B). The output is the bitvector representation (*VP*^*O*^, *VN*^*O*^, 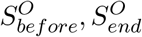) of a column *S*_*O*_ such that its values are the minimum of the two columns represented by the input bitvectors, that is, ∀*i* ∈ {0,1,…, *w* − 1} : 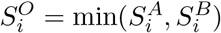 (step C).

First, we need to find *difference masks M*_*A*>*B*_ and *M*_*B*>*A*_, which describe cells where the score of *A* is higher than *B* and vice versa (step D). To do this, we first verify that the score differences *S*^*A*^ − *S*^*B*^ are in the range (−2*w*, 2*w*) (lines 20-25). We need the *popcount* operation for this; in most processors an *O*(1) specialized instruction exists, otherwise it can be calculated in *O*(log *w*) with normal arithmetic operations. We define a variable *D* split in *chunks*, where each chunk represents the score difference of *S*_*A*_ and *S*_*B*_ at a certain index. Each chunk is represented by log_2_ *w* + 2 bits. In this way, each chunk can accommodate a score difference, which takes a value between –2*w* and 2*w*. The chunks of D are initialized to point at the bit before the start of the chunk (step F, lines 26-26). Then, using *VP*^*A*^, *VP*^*B*^, *VN*^*A*^ and *VN^B^*, *D* is updated to point at the next bit and *M*_*A*>*B*_ and *M*_*B*>*A*_ are updated in parallel (step G, lines 32-37). This is repeated *O*(log *w*) times (step I, lines 31-37). Initializing the variable D takes *O*(log log *w*) time due to the chunk popcounts in line 5. Each iteration of the parallel update takes *O*(1) time, and there are *O*(log *w*) iterations. The runtime of calculating the difference) masks is therefore *O*(log *w*).

**Figure 3:**
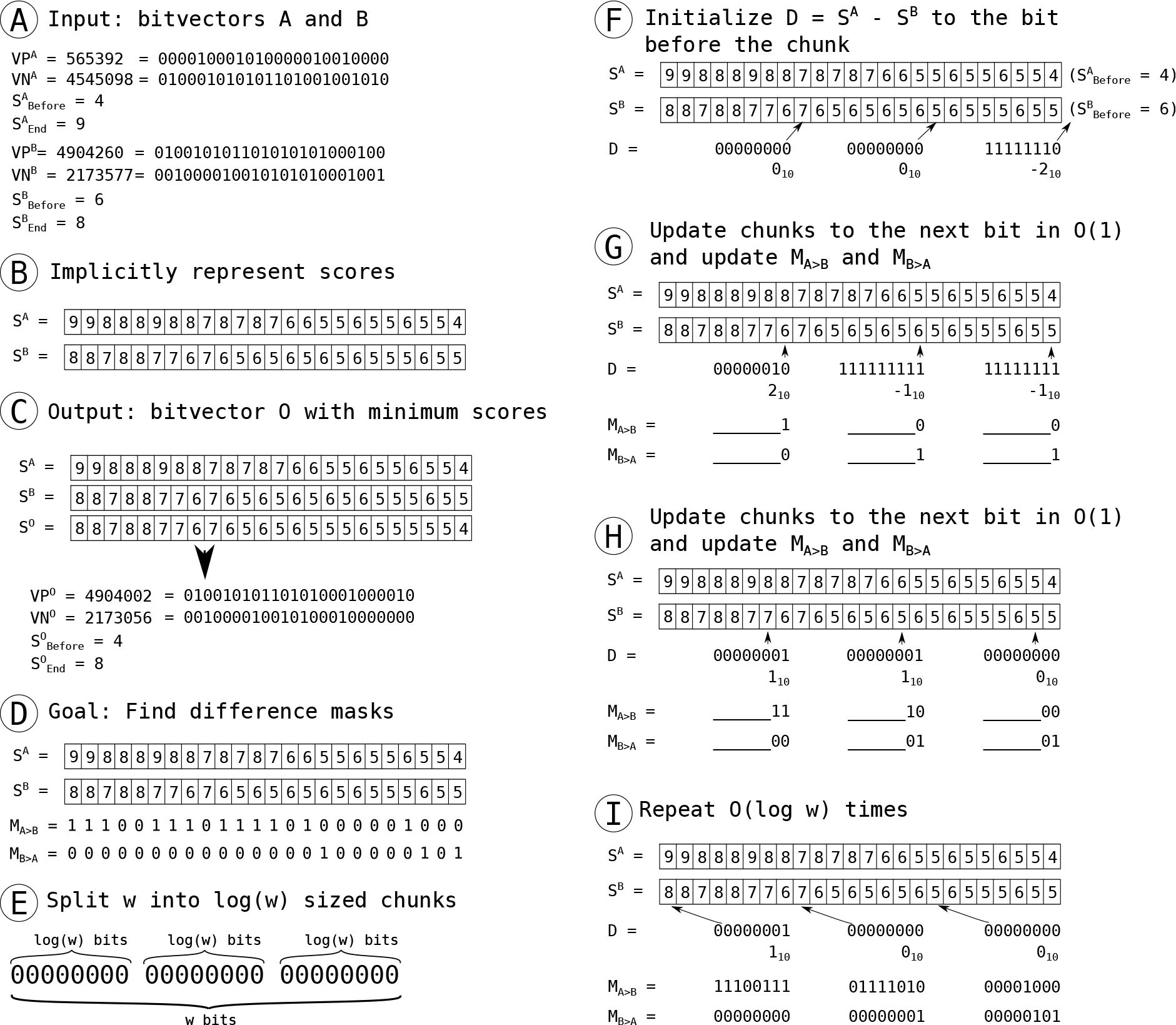
Overview of the bitvector merging algorithm

Once we have the difference masks *M*_*A*>*B*_ and *M*_*B*>*A*_, we can use them to merge the bitvectors. We first calculate a picking mask *M*_*p*_ which determines whether a bit needs to be taken from *S^A^* (line 41). Then we pick the values for *VP*^*O*^ and *VN*^*O*^ based on the picking mask. This gives the correct values for indexes where one of the bitvectors is greater or equal to the other in both the current and the previous index. But we still need to handle the case where one bitvector is greater at the current index and the other at the previous index. We do this by reducing the *VN*^*A*^ and *VN*^*B*^ vectors such that in those indices the output bitvector’s value cannot decrease (lines 42-43). Merging bitvectors, given a difference mask, is *O*(1). The total runtime is therefore *O*(log *w*).

However, in practice it is faster to precompute the difference masks for each possible 8-bit combination of 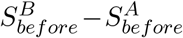, *VP*^*A*^, *VP*^*B*^, *VN*^*A*^ and *VN*^*B*^. Now, finding the difference masks takes 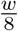 memory lookups. With the lookup, merging is *O*(*w*) but faster in practice than the *O*(log *w*) algorithm.

### 5.2 Changed minimum value algorithm

The *changed minimum value* of two bitvectors *old* and *new* is the minimum value at indices where the new bitvector has a smaller value than the old, ie. 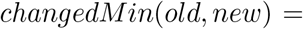 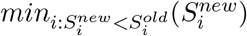. The changed minimum value can be calculated in *O*(log *w*) time by splitting the bitvector into chunks and calculating the value at each log *w*’th position in parallel, similarly to the difference mask algorithm. However, in practice it is faster to calculate the difference mask *M*_*new*<*old*_ and find all local minima where 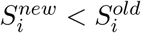 by using the *VP*, *VN* and *M*_*new*<*old*_ vectors. An index is a local minimum if *VP* is set to its left and *VN* is set either to its right or at the index. Then, each local minimum is processed one at a time. The score at the index is calculated using the definition of the implied scores *S*_*i*_ = *popcount*(*VP* _0․.*i*_) − *popcount*(*VN* _0․.*i*_). This takes *O*(*w*) time but in practice there are very few local minima, leading to a speedup over the *O*(log *w*) algorithm.

## 6 Experiments

We implemented the sequence-to-graph bitvector algorithm described here and the cell-by-cell algorithm by Navarro [14]. We created four kinds of graphs based on the *E. Coli* reference genome’s 10 000 first base pairs. Then, we simulated reads with 20x coverage from the reference using PBSIM [16] and aligned them to the graphs using both approaches. PBSIM produced 65 reads with an average length of 3kbp. The source code of the experiment is available at https://github.com/maickrau/GraphAligner/tree/WabiExperiments.

**Figure 4:**
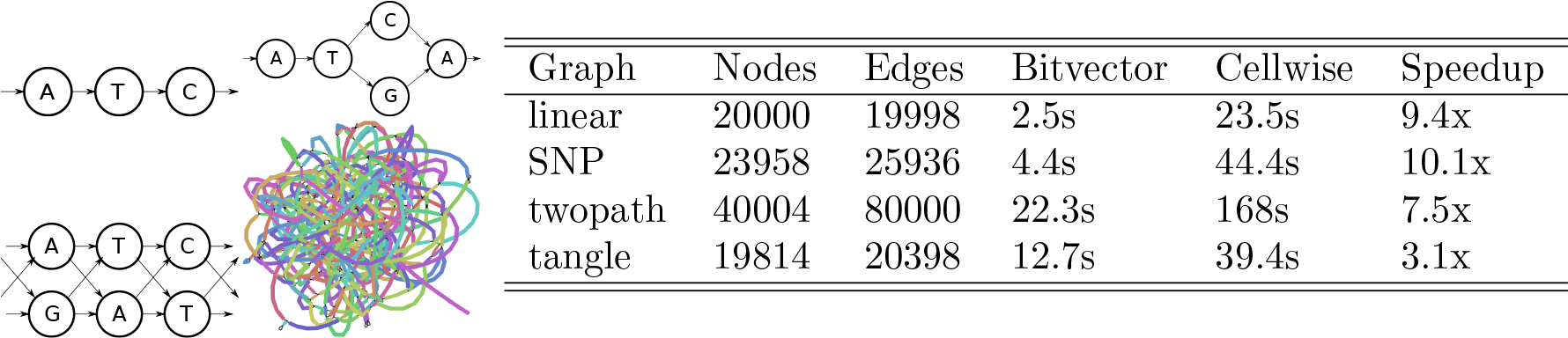
Left: Overview of the graphs used in the experiment. Left to right, top to bottom: Linear graph, SNP graph, twopath graph, tangle graph (visualized with Bandage [23]) Right: Summary of the runtime experiment.

Figure 4(left) shows the graph types used in the experiment. The first graph, the *linear graph,* is a linear chain of nodes. Aligning to this graph is equivalent to sequence-to-sequence alignment. The second graph, the *SNP graph*, is a linear chain of nodes with randomly inserted bubbles representing single nucleotide polymorphisms (SNPs). The SNPs are distributed at an average of one SNP per 10 base pairs. The third graph, the *twopath graph*, is an artificial worst case graph for the bitvector algorithm. Each node has two in-neighbors, which means that the *O*(log *w*) bitvector merging algorithm has to run for each node. For the first three graphs, neither algorithm’s runtime depends on the matched sequence, so the additional inserted nodes were given random labels. The fourth graph, the *tangle graph,* is a de Bruijn graph of the sequence with *k*=11. We chose *k* to be so small specifically to make the graph very cyclic and tangled. For each graph, we also included the reverse-complement strand to map reads simulated from the backwards strand, doubling the graph size from the original 10 000 base pairs and effectively mimicking a bidirectional graph.

Figure 4(right) shows a summary of the runtimes. Each number is an average of 10 runs. The bitvector approach is faster than the cell by cell approach in each graph. As expected from the time complexity, the difference is greater in the acyclic graphs. For the acyclic graphs, the bitvector algorithm achieves an about tenfold speed improvement. For the cyclic graph, the speedup is about threefold, suggesting that cycles are recalculated on average three times (linear speedup divided by cyclic speedup) instead of the theoretical worst case of 2*w* times.

## 7 Discussion

In this paper, we generalized two sequence-to-sequence algorithms to sequence-to-graph algorithms. For the Shift-And algorithm, the runtime for acyclic graphs matches the runtime of the linear version, and the runtime for cyclic graphs matches cell-by-cell comparison algorithms for graphs. For the bitvector alignment algorithm, the runtime includes an extra log *w* term due to the complexity of merging bitvectors and finding the changed minimum value. Despite the graph-based bitvector alignment algorithm’s higher worst case time complexity compared to previous cell-by-cell alignment algorithms, it still achieves a three-to tenfold speedup over cell-by-cell algorithms depending on the shape of the graph. Should an algorithm for merging bitvectors and finding the changed minimum score in *O*(1) time exist, that would lead to the bitvector graph algorithm being theoretically faster than cell-by-cell algorithms as well. The bitvector algorithm described here provides a basis for a practical algorithm for fast sequence-to-graph alignment.

## Acknowledgments

We thank Gonzalo Navarro for fruitful discussions on pattern matching on graphs. We are grateful for Dagstuhl Seminar 16351 on “Next Generation Sequencing - Algorithms, and Software For Biomedical Applications”, which sparked the idea to pursue this topic.

## A Appendix

## A.1 Proof of Theorem 1 (Vertical property)

*Proof*. It is clear from Recurrence 1 that *C*_*i,j*_ − *C*_*i,j-1*_ ≤ 1. Next, we have to prove the bound *C*_*i,j*_ − *C*_*i,j-1*_ ≥ −1 or, equivalently, *C*_*i,j-1*_ ≤ *C*_*i,j*_ + 1. We distinguish three cases, based on which of the three terms terms in Recurrence 1 takes a minimum (note that more than one term can have the minimum value).

*Case 1 (vertical*, *C*_*i,j*_ = *C*_*i,j-1*_ + 1). This case immediately implies *C*_*i,j*_ − *C*_*i,j-1*_ ≥ –1.

*Case 2 (diagonal*, *C*_*i,j*_ = *C*_*k*,*j-1*_ + Δ_*i,j*_ for 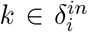). By Recurrence (1), we have *C*_*i,j−1*_ ≤ *C*_*k,j−1*_ + 1. Therefore, *C*_*i,j−1*_ ≤ *C*_*k*,*j−1*_ + 1 = *C*_*i,j*_ − Δ_*i,j*_ + 1 by the assumption of Case 2. It follows *C*_*i,j*_ ≥ *C*_*i,j−1*_ + Δ_*i,j*_ − 1, which implies *C*_*i,j*_ − *C*_*i,j−1*_ ≥ –1 since Δ_*i,j*_ ≥ 0 by definition.

*Case 3 (horizontal*, *C*_*i,j*_ = *C*_*k,j*_ + 1 *for* 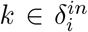). Our proof for this case is by induction on values of the cells in a row. That is, to prove the claim *C*_*i,j−1*_ ≤ *C*_*i,j*_ + 1 for cell *C*_*i,j,*_ we assume that it holds for all cells in the same row with smaller value, i.e. for all cells *C*_*i′,j*_ with *C*_*i′,j*_ < *C*_*i,j*_. Note that the claim holds for all cells with minimum value in each row, because they are covered by Case 1 or Case 2, but cannot fall under Case 3. By the induction hypothesis, we have *C*_*k,j−1*_ ≤ *C*_*k,j*_ + 1 and, by the assumption of Case 3, we get *C*_*k,j−1*_ ≤ *C*_*i,j*_. By Recurrence 1, we have *C*_*i,j−1*_ ≤ *C*_*k,j−1*_ + 1. Together, this implies that *C*_*i,j−1*_ ≤ *C*_*k,j−1*_ + 1 ≤ *C*_*i,j*_ + 1.

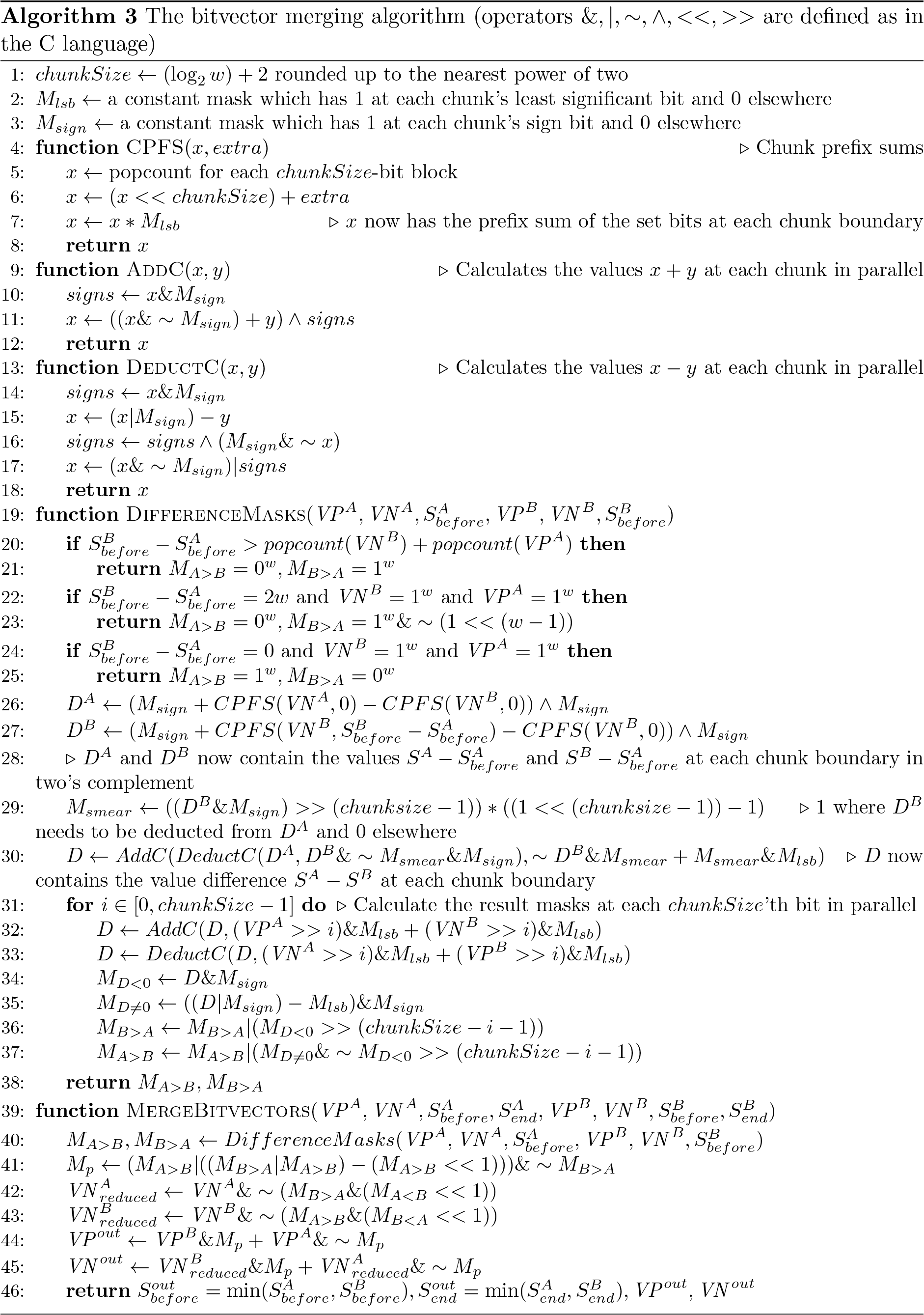

